# Heterologous synapsis in *C. elegans* is regulated by meiotic double-strand breaks and crossovers

**DOI:** 10.1101/2021.08.30.458290

**Authors:** Hanwenheng Liu, Spencer G. Gordon, Ofer Rog

## Abstract

Alignment of the parental chromosomes during meiotic prophase is key to the formation of genetic exchanges, or crossovers, and consequently to the successful production of gametes. In almost all studied organisms, alignment involves synapsis: the assembly of a conserved inter-chromosomal interface called the synaptonemal complex (SC). While the SC usually synapses homologous sequences, it can assemble between heterologous sequences. However, little is known about the regulation of heterologous synapsis. Here we study the dynamics of heterologous synapsis in the nematode *C. elegans*. We characterize two experimental scenarios: SC assembly onto a folded-back chromosome that cannot pair with its homologous partner; and synapsis of pseudo-homologs, a fusion chromosome partnering with an unfused chromosome half its size. We observed elevated levels of heterologous synapsis when the number of meiotic double-strand breaks or crossovers were reduced, indicating that the promiscuity of synapsis is regulated by break formation or repair. By manipulating the levels of breaks and crossovers, we infer both chromosome-specific and nucleus-wide regulation on heterologous synapsis. Finally, we identify differences between the two conditions, suggesting that attachment to the nuclear envelope plays a role in regulating heterologous synapsis.

## Introduction

Meiosis maintains the karyotypic stability of a species while generating genetic diversity through exchanges, or crossovers, between the parental genomes. Crossover formation, however, is inherently mutagenic and could disrupt genome integrity unless it integrates local sequence homology with chromosome contiguity. Failed crossover regulation—such as crossovers between homologous sequences that are on different chromosomes—leads to aneuploid gametes, causing inviability, sterility or congenital conditions in the offspring.

The synaptonemal complex (SC) is a conserved chromosomal interface that plays key roles in regulating crossovers and safeguarding a successful meiosis (Moses 1956, 1958; Fawcett 1956; Page and Hawley 2004; Rog and Dernburg 2013). The SC assembles between (or synapses) the parental homologous chromosomes (homologs) during meiotic prophase, bringing them into close juxtaposition and aligning them from end to end. In *C. elegans*, homologs are first brought into close proximity through specialized regions on each chromosome called Pairing Centers (Villeneuve 1994; MacQueen et al. 2005), which attach to the nuclear envelope and interact with cognate protein components (Phillips et al. 2005; Phillips and Dernburg 2006; Penkner et al. 2007; Sato et al. 2009). The SC starts assembling between the homologs at Pairing Centers (MacQueen et al. 2005; Rog and Dernburg 2015), stabilizing pairing interactions and extending them along the length of the chromosome (MacQueen et al. 2002). The physical tethering of the homologs helps ensure recombination occurs between homologous sequences on syntenic locations (Goldman and Lichten 2000).

The SC also ensures crossovers occur only between homologs by regulating crossover-promoting factors. Upon meiotic entry, many double-strand breaks (DSBs) are introduced (Keeney et al. 1997, Dernburg et al. 1998), a subset of which is repaired as crossovers. Crossover formation requires engagement with a homologous repair template, as well as recruitment of a suite of crossover-promoting factors to inter-homolog interface by the SC (Woglar and Villeneuve 2018; Li et al. 2018; Cahoon et al. 2019).

In spite of the ways in which the SC safeguards meiosis, its mode of assembly also leaves meiosis vulnerable to illegitimate crossovers. The extension of synapsis along the chromosome is sequence-independent and relies on contiguity (MacQueen et al. 2005). This property allows the SC to traverse small regions of heterozygosity, such as insertions and deletions, while maintaining overall register between homologs and in that way allowing crossovers to form (Hammarlund et al. 2005). However, the capacity of the SC for heterologous synapsis carries the risk of bringing together non-homologous chromosomes, thereby promoting illegitimate crossovers.

Two conditions that result in large-scale, cytologically-discernable heterologous synapsis are fold-back synapsis and synaptic adjustment. Fold-back synapsis occurs when a chromosome that fails to find its partner heterologously synapses its left and right halves. While unpaired X chromosomes remain mostly asynapsed in hermaphrodites (Phillips et al. 2005), fold-back synapsis is prevalent upon deletion of all Pairing Center proteins (Harper et al. 2011), likely due to the abundance of unassembled SC subunits. SC associated with unpaired chromosomes was also documented in X chromosome triploids (Mlynarczyk-Evans et al. 2013), and in as much as a third of nuclei in *C. elegans* males, which harbor a single, partner-less X chromosome (Checchi et al. 2014). Chromosomes can also undergo synaptic adjustment: a post-SC-assembly response to heterozygous translocations, inversions, deletions, insertions or fusions, which has been documented in many eukaryotes (Moses and Poorman 1981; Moses et al. 1982; Zickler and Kleckner 1999). Following initial synapsis based on local sequence homology, asynapsed overhangs, junctions and loops are minimized at the expense of heterologous synapsis.

Despite the prevalence of heterologous synapsis, and its potentially deleterious consequences, its regulation is poorly understood. A likely mode of regulation is through modulation of SC dynamics. Contrary to what its highly-ordered appearance in electron micrographs might suggest (Colaiácovo et al. 2003; Page and Hawley 2004), the SC displays liquid-like properties (Rog et al. 2017). SC fluidity is modulated in response to meiotic progression, and specifically to the repair of DSBs and formation of crossovers: upon crossover formation, the SC transitions into a less dynamic state (Pattabiraman et al. 2017). This effect is occurring at least partly *in cis* and can differentiate homolog pairs with and without crossovers (Machovina et al. 2016). However, whether DSBs or crossovers regulate heterologous synapsis has so far not been addressed.

Here we quantified heterologous synapsis in *C. elegans* worms carrying chromosomes that can undergo fold-back synapsis and synaptic adjustment. We tested the effect of genetic perturbations that alter the levels of DSBs or crossovers, and found increased heterologous synapsis in these two conditions. We calculated the rate of fold-back synapsis for unpaired chromosomes, and the effect of fusing an unpaired chromosome to paired homologs. Finally, by relying on the spatio-temporal organization of the *C. elegans* gonad, we document the dynamics of heterologous synapsis.

## Results

### Analyzing fold-back synapsis in *C. elegans*

To quantify the extent of fold-back synapsis, we used worms lacking HIM-8, a protein necessary for pairing, and subsequent synapsis, of the X chromosomes (Phillips et al. 2005). Consequently, the DSBs that are formed on the unpaired X chromosomes are repaired without forming crossovers.

We inferred fold-back synapsis from the absence of asynapsed X chromosomes. We labeled the chromosomal axis protein HTP-3 (Goodyer et al. 2008) and the SC subunits SYP-1 (MacQueen et al. 2002) or SYP-5 (Hurlock et al. 2020; Zhang et al. 2020). Initial synapsis yields overlapping axis and SC staining for the autosomes, while the two X chromosomes are easily distinguished as asynapsed: axial structures that lack SC staining (Fig 1a, left and Fig 1b, top). If either or both X chromosomes are folded back and self-synapsed, we would see one or zero asynapsed regions, respectively (Fig 1b, middle and bottom). The progression of fold-back synapsis was easy to assess in the *C. elegans* gonad due to its spatial-temporal organization. We quantitated progression by dividing the pachytene region of the gonad, when crossovers mature, into three bins of equal length, roughly spanning early-, mid- and late-pachytene (Fig S1; see Methods; Schedl 1997).

**Fig. 1.**
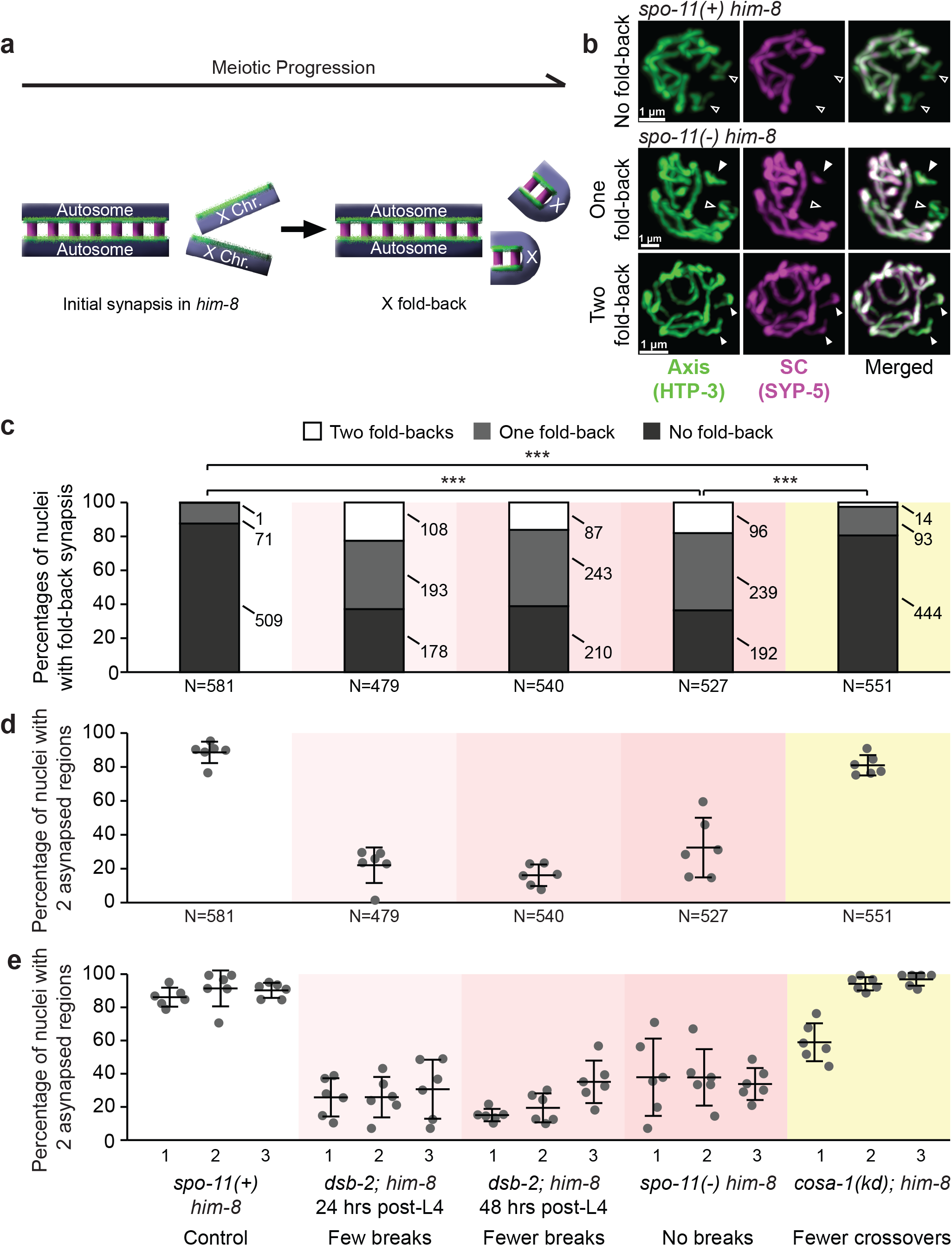
DSBs and crossovers affect fold-back synapsis. a. Schematic of the experimental systems to assess fold-back synapsis in standalone X chromosomes. Initially, only the autosomes synapse, while pairing and synapsis of the X chromosomes is prevented by the absence of HIM-8. As meiosis progresses, some of the unpaired X chromosomes fold back and self-synapse (in the case shown, both X chromosomes). b. Co-immunofluorescence shows asynapsed regions. Top row: a nucleus from *spo-11(+) him-8* lacking fold-back. Two asynapsed regions (axis staining unaccompanied by SC staining) can be seen, indicating no fold-back synapsis. Middle and bottom row: nuclei with no DSBs from *spo-11(−) him-8* with one or no asynapsed region visible (middle and bottom rows, respectively), indicating fold-back synapsis of one or both unpaired X chromosomes. Green, HTP-3 (axis). Magenta, SYP-1 (SC). Hollow arrowheads, X chromosomes. White arrowheads, fold-back synapsis of X chromosomes. Scale bars=1 μm. c. Percentage of nuclei with two, one, or zero X chromosomes that underwent fold-back synapsis in each strain. Numbers of nuclei in each category are shown on the right of each stack. Fold-back synapsis is rare in *spo-11(+) him-8*, prevalent in the absence of DSBs, and uncommon in the absence of crossovers (all pairwise comparisons p<0.0005, Pearson’s chi-squared test). N represents total number of nuclei analyzed. Relevant genotypes and their effects are indicated at the bottom. d. Percentage of nuclei with two asynapsed regions, indicating fold-back synapsis on both unpaired X chromosomes. Number of nuclei with fold-back synapsis increases sharply upon perturbation of DSBs, but only slightly when crossover level is lowered (p=0.05764, Student’s t-test). N represents the number of nuclei from the 10 gonads assessed for each genotype. Each dot represents one gonad. Bars show mean ± SD. Relevant genotypes and their effects are indicated at the bottom. e. Dynamics of fold-back synapsis are affected by the levels of DSBs and crossovers. Perturbations of DSBs raised the amount of fold-back synapsis, but did not reveal significant trends throughout pachytene. Bars show mean ± standard deviation (SD). N=10 gonads for each bin. Relevant genotypes and their effects are indicated at the bottom.

We first analyzed fold-back synapsis in *spo-11∷aid him-8* animals grown on NGM plates (referred to throughout as *spo-11(+)*), which do not exhibit any perturbation to meiotic progression, specifically with regards to the formation of DSBs and crossovers (Zhang et al. 2018; Almanzar et al. 2021), and therefore serve as our control. Consistent with previous analyses (Phillips et al. 2005), without HIM-8 only a minimal number of X chromosomes were synapsed: 71 of 581 nuclei contained one folded-back X chromosome and only one of 581 nuclei had both X chromosomes folded-back (Fig 1c).

### DSBs inhibit fold-back synapsis

DSB and crossover formation are tightly regulated by multiple surveillance mechanisms that impact chromatin and the SC in both chromosome-specific and nuclear-wide fashion (Rosu et al. 2013; Stamper et al. 2013; Machovina et al. 2016; Pattabiraman et al. 2017). We wondered whether the same surveillance mechanisms might also regulate heterologous synapsis.

We analyzed the effects of DSBs on fold-back synapsis by comparing three conditions with progressively reduced DSB number. Worms lacking the DSB-promoting factor DSB-2 exhibit progressive reduction in DSB number with age (Rosu et al. 2013), resulting in an average of 0.92 and 0.31 DSBs per chromosome pair in worms 24 and 48 hours post-L4 stage, respectively (see Methods for calculations). These numbers are dramatically lower than the estimated 2-5 DSBs per homolog pair in wild-type animals (Colaiácovo et al. 2003; Mets and Meyer 2009; Rosu et al. 2013; Woglar and Villeneuve 2018), and entails that many chromosomes in *dsb-2* animals do not receive any DSBs. To eliminate DSBs altogether we degraded SPO-11, which catalyzes meiotic DSBs (Keeney et al. 1997; Dernburg et al. 1998), using the auxin-degradable allele *spo-11∷aid* (Zhang et al. 2015, 2018; Almanzar et al. 2021). When raised on auxin plates, *spo-11∷aid* worms lacked DSBs and crossovers altogether, and are referred to hereon as *spo-11(−)*.

*dsb-2* worms at 24 hours post-L4 exhibited an increase of nuclei with one or zero asynapsed regions compared to *spo-11(+)* controls (Fig 1c, one folded-back chromosomes: 193 out of 479; two folded-back chromosomes: 178 out of 479; versus *spo-11(+)*: p<0.0005, Pearson’s chi-squared test). Fold-back synapsis was also elevated in *dsb-2*, 48 hours post-L4 (Fig 1c; 243 one-fold-back and 87 two-fold-back chromosomes out 540 total) and in *spo-11(−)* (Fig 1c; 239 one-fold-back and 96 two-fold-backs out of 527). While the increased rate of synapsis upon DSB perturbation could have potentially been caused by homologous synapsis of the X chromosomes, we find this unlikely. First, we observe many nuclei with only one asynapsed region (Fig 1c). Second, closer examination of nuclei with no asynapsed regions found most of them to have 7 rather than 6 SC threads (29/32 nuclei with zero asynapsed regions had 7 SC threads in *spo-11(−) him-8* worms), consistent with two fold-back synapsis events.

Taken together, we find that DSBs are crucial for suppressing fold-back synapsis. Furthermore, the similar increase of fold-back synapsis in *spo-11(−)* (no breaks) and *dsb-2*, 24 hours (0.46 DSBs per X chromosome) suggests that the mechanism underlying the suppression of fold-back synapsis does not rely on a per-chromosome inhibition but rather on a nucleus-wide effect.

### Crossovers inhibit fold-back synapsis

In *C. elegans*, an orchestrated response to chromosomes lacking crossovers is thought to help ensure each chromosome receives a crossover (Rosu et al. 2013; Stamper et al. 2013; Machovina et al. 2016). Such a response will be activated in the mutant scenarios analyzed above since not enough DSBs are made to generate at least one crossover on each homolog pair. (This would take place in conjunction with the response to the unpaired X chromosomes (Harper et al. 2011).) Furthermore, the SC directly responds to crossover formation by locally expanding (Libuda et al. 2013; Woglar and Villeneuve 2018). We therefore examined the role of crossovers in regulating fold-back synapsis.

COSA-1 is essential for the maturation of DSBs to crossovers (Yokoo et al. 2012). We attached *aid* degron to the N terminus of *cosa-1* using CRISPR/Cas9 (Fig S2; Zhang et al. 2015). Our initial construct—*aid∷cosa-1* with the plant TIR-1 E3 ligase driven by the *sun-1* promotor (*sun-1p∷tir-1*)—had no discernable effect on COSA-1 function in the absence of auxin, and 70% reduction in viable self-progeny in the presence of auxin (p=0.001438, Welch’s t-test, α=0.05, same below). This reduction in viable progeny did not reach the level of *spo-11(−)* (Fig S3a) or a *cosa-1* null mutant (Yokoo et al. 2012), suggesting that the degradation of AID∷COSA-1 was incomplete. Upon switching the TIR-1 promotor from *sun-1p* to *gld-1p* (Chen et al. 2020), we observed 87% reduction in viable self-progeny (Fig S3a, p=0.04753 *versus sun-1p*, p=0.0009 *versus gld-1p* on NGM, Welch’s t-test), suggesting it was more efficient in degrading AID∷COSA-1. We refer to these worms, when grown on auxin, as *cosa-1(kd)*, for COSA-1 <u>k</u>nock<u>d</u>own.

To quantify the number of crossovers in *cosa-1(kd)* worms, we quantified the number of joined homologs immediately prior to the first meiotic division (DAPI bodies). Wild-type animals form six DAPI bodies, one for each of the six homolog pairs. A failure to form crossovers results in 7-12 DAPI bodies. *cosa-1(kd); him-8* worms exhibited an average of 9.0 DAPI bodies, similar to *zim-2 zim3 him-8* triple mutant, where only 2 of the 6 homologs (chromosomes I and II) pair and form crossovers (Fig S3b and S3c; p=0.3976, Student’s t-test; Phillips and Dernburg 2006). *cosa-1(kd)* therefore reduces crossovers number so that only about 1/3 of the homolog pairs form a crossover.

We observed only a slight increase in fold-back synapsis in *cosa-1(kd)* worms relative to *spo-11(+)* animals, with 93 out of 551 nuclei that contained only one folded-back X chromosome, and 14 out of 551 nuclei that contained two folded-back X chromosome (Fig 1c; p=0.0001 *versus spo-11(+)*, Pearson’s chi-square test). Notably, *cosa-1(kd)* worms exhibited a dramatically different fold-back synapsis distribution from *dsb-2*, 24 hours post-L4 (p<0.0005, Pearson’s chi-squared test) despite similar levels of crossovers in these two conditions (3.0 and 3.6 crossovers per nucleus in *cosa-1(kd)* and *dsb-2*, 24 hours, respectively; Fig S3c; Rosu et al. 2013). Since *cosa*-*1(kd)* causes a reduction of crossovers despite high levels of DSBs (Yokoo et al. 2012) while in *dsb*-*2* both DSBs and crossovers are reduced (Rosu et al. 2013), the difference we observed suggests that it is DSBs and not crossovers that play a key role in suppressing fold-back synapsis.

To highlight the differences, we focused on the percentage of nuclei with two asynapsed regions, where no fold-back synapsis occurred (Fig 1d). This metric facilitated tracking of fold-back synapsis along meiotic progression. The percentage of nuclei with two asynapsed regions stayed at a high level in *spo-11(+)* throughout pachytene (Fig 1e; bin 1 to bin 3 averages, 85%, 91%, and 90%, respectively). These rates lowered by three quarters in both *dsb-2*, 24 and 48 hours post-L4, and by two-thirds in *spo-11(−)* (Fig 1e). While most conditions did not reveal significant trends throughout pachytene, an exception was *cosa-1(kd)*, where fold-back synapsis was higher in early pachytene, but decreased with meiotic progression (Fig 1e).

### DSBs and crossovers affect heterologous synapsis between pseudo-homologs

We next tested another condition resulting in heterologous synapsis: a pair of pseudo-homologs of unequal sizes. We constructed worm strains carrying a single copy of a fusion chromosome, *ypT27*, in which the left end of chromosome V is joined to the right end of the X chromosome (Lowden et al. 2008). We also deleted *him-8*, which prevented pairing of the two X chromosomes. In animals heterozygous for the fusion chromosome, the pseudo-homologs initially pair and synapse through chromosome V homology, allowing for crossover formation on that chromosome; the fused X chromosome overhangs at the terminus, and the standalone X chromosome is left unpaired and asynapsed (Fig 2a, left). Subsequently, the pseudo-homologs can be brought into full synapsis through various forms of heterologous synapsis. These included synaptic adjustment (Fig 2a, right, first and second rows), which has been documented in male worms carrying a similar pair of pseudo-homologs (Henzel et al. 2011); folding back of the fused X chromosome (Fig 2a, right, third row); or a combination of the two (Fig 2a, right, fourth row). We visualized heterologous synapsis in the pseudo-homologs as we did fold-back synapsis, and inferred heterologous synapsis from the absence of asynapsed regions, which would occur upon complete heterologous synapsis of the pseudo-homologs or by complete fold-back synapsis of the standalone X chromosome (Fig 2b).

**Fig. 2.**
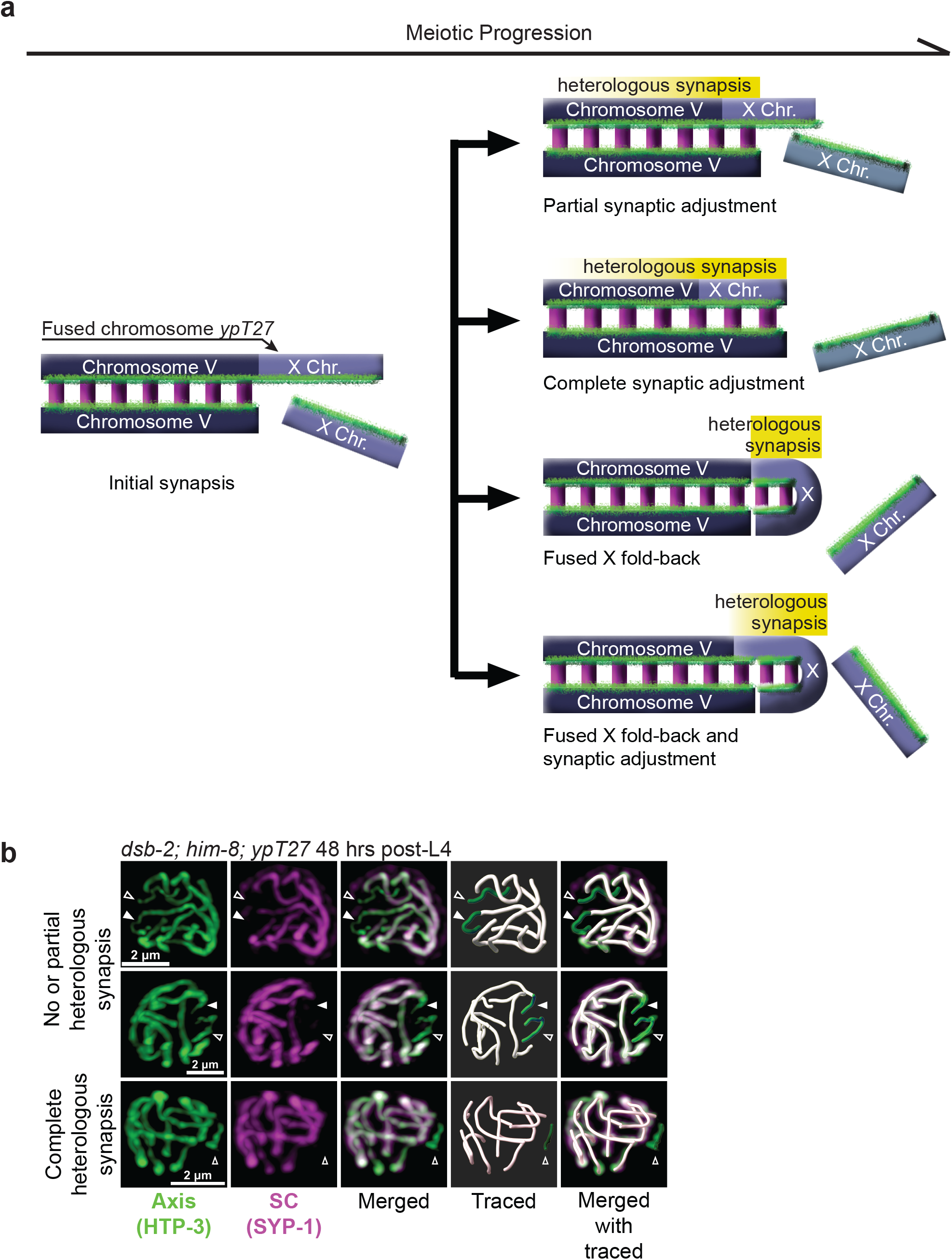
Heterologous synapsis of pseudo-homologs. a. Experimental system to detect outcomes of heterologous synapsis between a chromosome pair of unequal lengths. One of the chromosomes is a standalone chromosome V (dark blue), and the other is a fused chromosome of V and X (light blue). Left: initial synapsis between the fused chromosome and chromosome V leaves an X chromosome overhang. Right: as meiosis progresses, the two pseudo-homologs heterologously synapse. Partial synaptic adjustment (first row) leaves part of the overhang of the X chromosome. Complete synaptic adjustment (second row) means that the fused chromosome and the standalone chromosome V are adjusted to equal lengths. In both cases, much of the right side of chromosome V is synapsed with a heterologous partner (yellow shading). Fold-back synapsis (third row) and a combination of fold-back synapsis and synaptic adjustment (fourth row) can also happen on the fused X chromosome. b. Co-immunofluorescence shows asynapsed regions for heterologous synapsis in *ypT27+/−*; *dsb-2; him-8* worms, 48 hours post-L4. Top and middle rows: two asynapsed regions are visible in each nucleus, indicating no or partial heterologous synapsis. Bottom row: only one asynapsed region is visible, indicating complete heterologous synapsis. Green, HTP-3 (axis). Magenta, SYP-1 (SC). White skeletons in traced images, synapsed region. Green skeletons in traced images, asynapsed regions. Hollow arrowheads, standalone X chromosome. Filled arrowheads, X chromosome overhang on the fused chromosome. Scale bars=2 μm.

Once again, *spo-11(+)* served as our control. In a total of 778 nuclei we scored, a great majority—594 nuclei—contained two asynapsed regions, 165 and 19 nuclei had only one or zero asynapsed region, respectively (Fig 3a). This is in line with our fold-back synapsis data above, and with observations on a similar pair of pseudo-homologs in males (Henzel et al. 2011). This observation reiterated the notion that heterologous synapsis is uncommon when other meiotic processes are intact. Notably, however, we observed more heterologous synapsis in nuclei with the fusion chromosome compared to those with the two standalone chromosomes (Figs 1c and 3a; percentage of nuclei with two asynapsed regions, with:without fusion chromosome=73%:88%; p<0.0005, Pearson’s chi-squared test). Since the difference between these two experimental scenarios is the fusion of X chromosome on the pseudo-homolog, our data suggest a role for fusion to a synapsed chromosome in regulating heterologous synapsis.

**Fig. 3.**
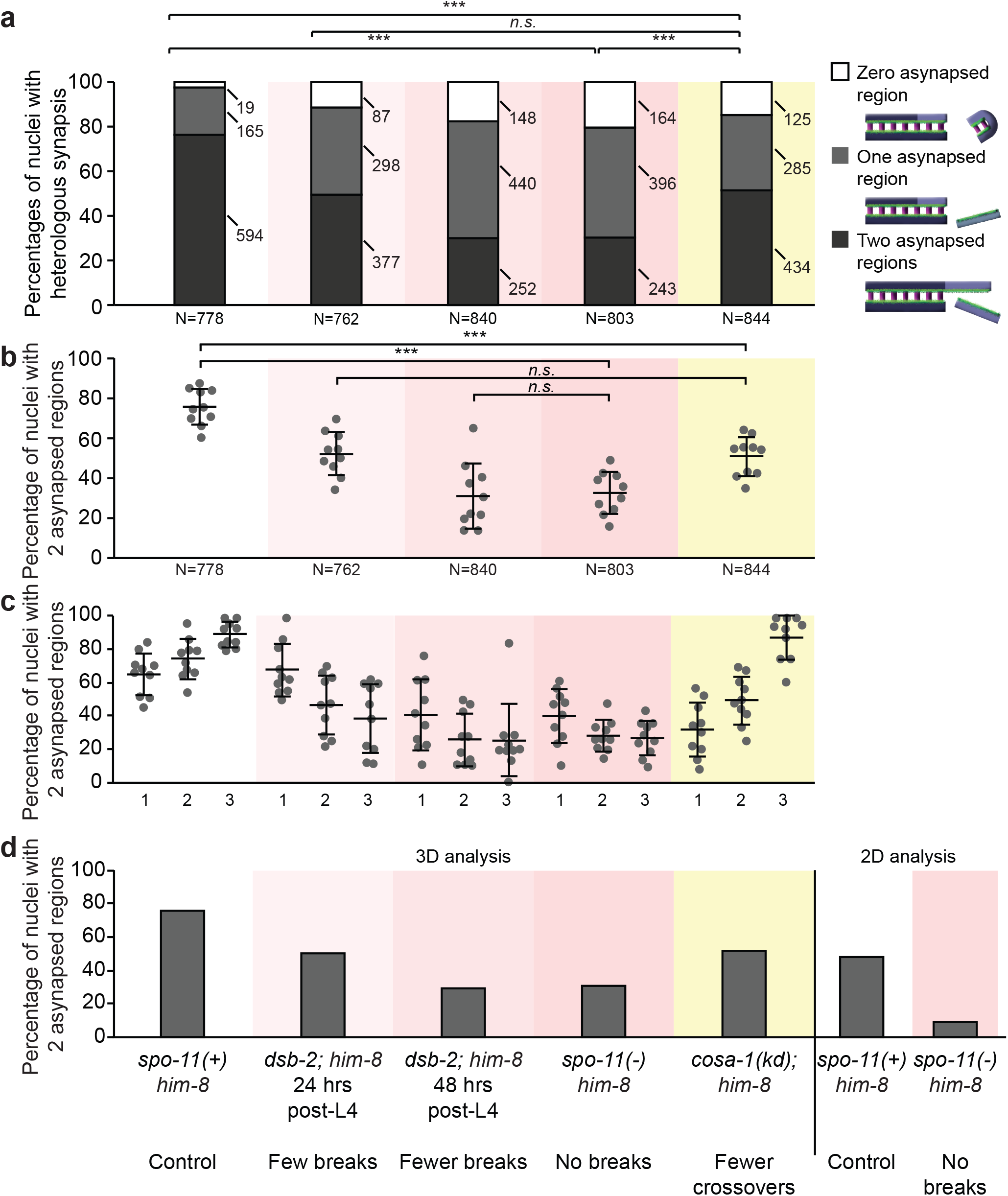
DSBs and crossovers affect heterologous synapsis in pseudo-homologs. a. Percentage of nuclei with zero, one, or two asynapsed regions in each strain. Numbers of nuclei in each category are shown on the right of each stack. Nuclei with zero or one asynapsed region (implying complete heterologous synapsis) are rare in *spo-11(+) him-8*, but common in the absence of double-strand breaks and crossovers (all pairwise comparisons p<0.0005, Pearson’s chi-squared test). N represents total number of nuclei analyzed. Relevant genotypes and their effects are indicated at the bottom. b. Heterologous synapsis is affected by the levels of DSBs and crossovers. Nuclei with two asynapsed regions decrease with lower levels of double-strand breaks or crossovers (p<0.0005 for all comparisons with *spo-11(+)*, Student’s t-test). N represents total number of nuclei from the 10 gonads assessed for each genotype. Each dot represents one gonad. Bars show mean ± SD. All worms contain *ypT27*/+ pseudo-homolog. Relevant genotypes and their effects are indicated at the bottom. c. Dynamics of heterologous synapsis are affected by the levels of DSBs and crossovers. Percentage of nuclei with two asynapsed regions, indicating partial or no heterologous synapsis, are shown. As DSBs are progressively eliminated, the levels of heterologous synapsis increase. Bars show mean ± standard deviation (SD). N=10 gonads for each bin. All worms contain *ypT27*/+ pseudo-homolog. Relevant genotypes and their effects are indicated at the bottom. d. The effects of eliminating DSBs are robust, regardless of the method of scoring synaptic adjustment. Each bar represents total nuclei scored. 3-dimensional (3D) analysis (left) was performed by analyzing reconstructions of confocal z-stacks. 2-dimensional (2D) analysis (right) was performed on maximum-intensity orthogonal projections of independent datasets. N values for 3D analysis are the same as in Fig 3B. N=427 (*spo-11(+)*) and N=458 (*spo-11(−)*) for 2D analysis. All worms contain *ypT27*/+ pseudo-homolog. Relevant genotypes and their effects are indicated at the bottom.

As we experimentally diminished the number of DSBs, asynapsis decreased. In *dsb-2*, 24 hours post-L4, around half of the total nuclei (377 out of 762) contained two asynapsed regions, and around 1/3 contained one asynapsed region (298 nuclei). In *dsb*-*2* animals at 48 hours post-L4, with fewer DSBs, most X chromosomes underwent heterologous synapsis (440 and 148 nuclei out of 840 with one and zero asynapsed regions, respectively; Fig 3a). This pattern also occurred in *spo-11(−)*, where no breaks take place (Fig 3a; p=0.29, *dsb-2*, 48 hours post-L4 *versus spo-11(−)*, Pearson’s chi-square test). The similarity between *dsb*-*2*, 48 hours post-L4 and *spo*-*11(−)* points to a role for a critical number of DSBs in suppression of heterologous synapsis. Reduced crossovers in *cosa-1(kd)* also increased heterologous synapsis, with 434 out of 844 nuclei containing two asynapsed regions, and 285 and 125 nuclei bearing one or zero asynapsed regions, respectively (Fig 3a; p<0.0005 *versus spo-11(+)*, Pearson’s chi-square test).

As above, we focused on the percentage of nuclei with two asynapsed regions to facilitate comparisons between the different conditions. As pachytene progressed, the percentage of nuclei with two asynapsed regions increased in our *spo-11(+)* control, from a 66% average in bin 1 to 75% in bin 2 to 90% in bin 3 (Fig 3b), indicating a decrease in heterologous synapsis. This came as a surprising contrast with previous observations in *C. elegans* males (Henzel et al. 2011) and in other systems (Moses and Poorman 1981; Moses et al. 1982; Bojko 1990; Torgasheva et al. 2013), where heterologous synapsis increased with meiotic progression. Interestingly, the dynamics of heterologous synapsis differed between perturbations of DSBs and crossovers: while the trend of heterologous synapsis in *cosa-1(kd)* mirrored that of the *spo-11(+)* control, it was reversed in the three conditions of perturbed DSBs, exhibiting increased heterologous synapsis with meiotic progression (Fig 3c). These opposite dynamics suggest that the underlying mechanism of heterologous synapsis suppression by DSBs and crossovers might not be the same.

Finally, to assess the precision of our methodology, we also analyzed asynapsed regions using maximum-intensity projections (2D analysis). This approach allows for a higher throughput while compromising precision. 2D analysis led to an apparent decrease in nuclei with two asynapsed regions in *spo-11(+)* (Fig 3d; two asynapsed regions, 2D *versus* 3D=49% *versus* 76%). This is likely because some asynapsed regions were obstructed by overlapping chromosomes. Nonetheless, when comparing *spo-11(+)* and *spo-11(−)* worms we observed significantly fewer nuclei with two asynapsed regions in *spo-11(−)* worms (9%; p<0.005 compared to *spo-11(+)*, Pearson’s chi-squared test), recapitulating the trend we observed in the 3-dimensional analysis above.

### Calculating heterologous synapsis rates for the standalone and fused X chromosomes

We used the distribution of nuclei with different numbers of asynapsed region to estimate the rate of complete fold-back synapsis for an unpaired X chromosome, R_S_ (for <u>s</u>tandalone; Fig 1c), and the rate of complete heterologous synapsis for a fused X chromosome, R_F_ (for <u>f</u>used; Fig 3a). First, we fitted the fold-back synapsis dataset for each condition to a binomial distribution and solved for the best-fit R_S_ (Fig 4; see Methods). In most conditions, we observed a good fit to a binomial distribution (e.g., p=0.66029 in *spo-11(+)*, Chi-squared goodness-of-fit test), suggesting that, within each nucleus, the two asynapsed chromosomes undergo heterologous synapsis independently. Subsequently, we calculated R_F_ for the pseudo-homologs: we used our calculated R_S_—assuming it describes well the behavior of the standalone X—to solve for the best-fit R_F_ based on the distribution of asynapsed regions in the pseudo-homolog datasets (Fig 4). These, too, exhibited mostly good fits to binomial distribution with two different rates.

**Fig. 4.**
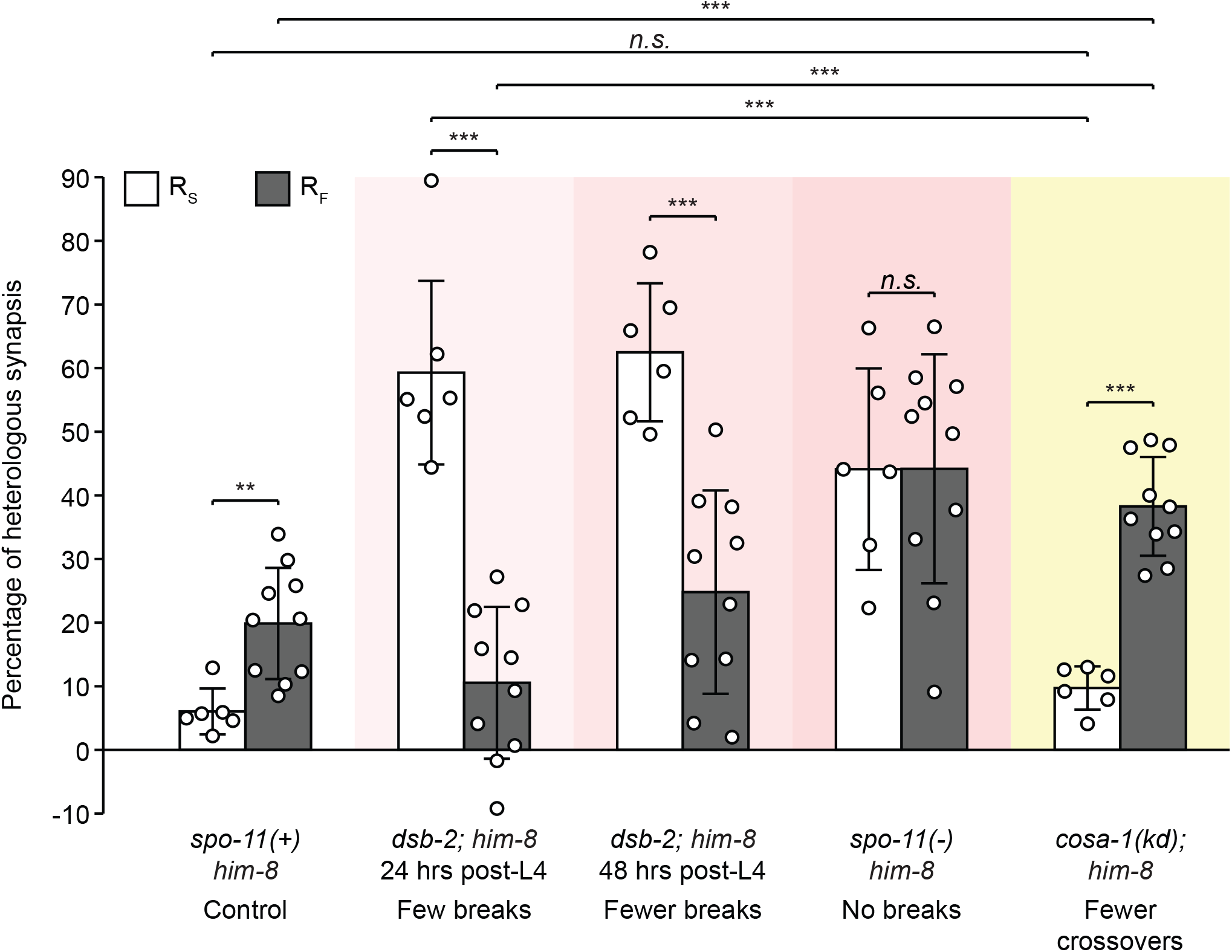
Heterologous synapsis rates for standalone and fused X chromosomes. R_S_ and R_F_ in the different experimental conditions. In *spo-11(+)* and *cosa-1(kd)*, R_F_ is larger than R_S_. Perturbation of DSBs causes generally larger R_S_ and R_F_ values, but reverses the trend: minimally reducing DSB mostly affect R_F_, but the two values are equalized upon complete depletion of DSBs. Bars show average R_F_ and R_S_. N=6 gonads per condition for R_S_. N=10 gonads per condition for R_F_, represented by dots. Error bars show mean ± SD. Relevant genotypes and their effects are indicated at the bottom.

With these rates in hand, we directly compared the effect of fusion on complete heterologous synapsis. In *spo-11(+)* controls, both R_S_ and R_F_ remained relatively low. However, the rate of complete heterologous synapsis was three-fold higher when fused to an autosome rather than standalone (Fig 4; average R_S_=6.1%, R_F_=19.9; p=0.0007, Welch’s t-test). The same trend was observed again in *cosa-1(kd)*, although both rates were correspondingly higher (Fig 4; compared to *spo-11(+)*, R_S_=9.7%, p=0.0987, Student’s t-test; R_F_=38.3%; p<0.0005, Student’s t-test). Interestingly, the trend reversed when DSBs were perturbed. The standalone X chromosome was more likely to engage in heterologous synapsis than the fused X chromosome. As DSBs were gradually depleted, the gap between R_S_ and R_F_ gradually diminishes (Fig 4).

## Discussion

In this study, we perturbed the levels of DSBs and crossovers to examine their effect on heterologous synapsis. We found that heterologous synapsis is an uncommon event in nuclei with unperturbed DSB and crossover levels. However, decreased number of DSBs increased heterologous synapsis and also changed its dynamics, causing more heterologous synapsis as meiosis progressed. Decreasing crossover number also increased heterologous synapsis, but to a lesser extent and without affecting synapsis dynamics. Finally, by comparing worms with two different karyotypes, we were able to deduce how attachment to a synapsed chromosome, and through it to the nuclear envelope, regulates heterologous synapsis.

### Heterologous synapsis is uncommon

Successful meiosis maintains karyotypic stability by initiating synapsis using homology-based mechanisms. Across organisms, these include both DSB-dependent and -independent mechanisms (Zickler and Kleckner 1999). Worms utilize one of the best characterized DSB-independent mechanisms, where synapsis initiation relies on interactions between Pairing Centers (Villeneuve 1994; MacQueen et al. 2005), attachment to the nuclear envelope (Penkner et al. 2007), and force generation by dynein (Sato et al. 2009). Consistent with the robustness of these mechanisms, we find low levels of heterologous synapsis in worms lacking the X chromosome Pairing Center protein HIM-8, as long as DSBs and crossovers are unperturbed (Fig 1). In worms where the unpaired X chromosome was fused to a synapsed autosome, the heterologous synapsis rate was significantly higher but still representing a small subset of chromosomes (Figs 2 and 3). Since the SC helps recruit crossover-promoting factors (Woglar and Villeneuve 2018; Li et al. 2018; Cahoon et al. 2019), such suppression of heterologous synapsis helps limit illegitimate recombination: exchanges between homologous sequences that are not syntenic.

Interestingly, our data shows that heterologous synapsis does not increase with meiotic progression. We observed relatively constant levels of fold-back synapsis of the unpaired X chromosomes (Fig 1e) and decreasing levels of heterologous synapsis for worms with a fused X chromosome (Fig 3c). These findings contrast with earlier studies of synaptic adjustment in *C. elegans* males (Henzel et al. 2011), and in other systems (Moses and Poorman 1981; Moses et al. 1982; Bojko 1990; Torgasheva et al. 2013), which observed progressively more adjustment with meiotic progression. One potential explanation for the apparent discrepancy is that we conflated heterologous synapsis with desynapsis, which occurs at the end of pachytene (MacQueen et al. 2002) and is particularly prevalent on chromosomes lacking crossovers (Machovina et al. 2016). This is unlikely, since our region of interest in the gonads ended prior to regions of prevalent desynapsis. Also, the gradual decrease in heterologous synapsis started already in bin 2 (Figs 1e & 3c), arguing that late-pachytene events are not solely responsible for our observations.

The propensity of the SC to synapse the chromosome from end to end, as well as the abovementioned studies, have led to the suggestion that synapsed chromosomes represent a low energy state. However, our data suggests that with meiotic progression, the SC has a smaller propensity for heterologous synapsis, potentially representing a proofreading-like mechanism. Interestingly, association of SC proteins with the lone X chromosome in males has been observed specifically in early pachytene (Checchi et al. 2014). The decreasing propensity for heterologous synapsis could be dependent on modulated biophysical properties of the SC, as previously suggested (Pattabiraman et al. 2017). In addition, loss of heterologous synapsis in mid-pachytene—a meiotic stage where homologous synapsis is unaffected—suggests a role for sequence homology or a process that depends on homology (such as repair intermediates) in stabilizing the SC specifically between homologous chromosomes.

In mutants with reduced levels of DSBs, but not with reduced crossovers, the dynamics of heterologous synapsis for the fused X is reversed, with increased heterologous synapsis with meiotic progression (Fig 3c). This result points to a role for repair intermediates in regulating the SC. The similar dynamics between conditions with reduced DSBs (0.3-0.9 per chromosome) and complete lack of crossovers suggests that it is not the presence of a DSB on a particular chromosome that restricts heterologous synapsis, but rather the global level of DSBs that transitions the SC to a restrictive state.

### Different effects of breaks and crossovers on heterologous synapsis

A major finding of our work is that almost all conditions where DSB or crossover levels are perturbed resulted in increased heterologous synapsis. In addition, comparing these conditions is telling.

While both *dsb-2*, 24 hours post-L4 and *cosa-1(kd)* animals form a similar number of crossovers—2-3 per nucleus, or ~40% of chromosomes undergoing a crossover—the former exhibited increased fold-back synapsis (Figs 1 and 4). Since the main difference between the two conditions are DSB levels (0.9 per chromosome *versus* unperturbed levels, likely 2-5 DSBs per chromosome), our data suggests that DSBs or repair intermediates suppress fold-back synapsis on the unpaired X chromosomes. Strong effect for DSBs compared to crossover on fold-back synapsis makes mechanistic sense as well, considering that the unpaired X chromosomes do not undergo crossovers (Phillips et al. 2005).

We observed similar increases in fold-back synapsis among all three conditions with reduced DSBs, regardless of their severity (Fig 1c-d). This suggests that DSBs affect fold-back through a nucleus-wide mechanism that requires a threshold of DSBs, and not through a direct effect on the folded-back chromosome. If the latter were the case, we would have expected a gradual effect with decreasing DSBs.

Crossovers also play a role in suppressing heterologous synapsis. This is particularly apparent on the fused X chromosome, where reduction of crossovers leads to doubling of R_F_ from 19% to 38%. While our methodology does not allow us to distinguish between different modes of heterologous synapsis—i.e. fold-back synapsis *versus* synaptic adjustment—an attractive possibility is that crossovers, by virtue of linking the homologs, impose a physical barrier that prevents complete synaptic adjustment. Consistent with a *cis* model for suppression is the dose-dependent response of the fused X to reduction in DSBs (Fig 4), which entails a gradual decrease in the chance the homolog pair undergoes a crossover. This result is also consistent with the results of Henzel et al. (2011) in *C. elegans* males, where despite significant synaptic adjustment, complete adjustment of the pseudo-homologs is rarely achieved; and with the results of Torgasheva et al. (2013), which identified a role for crossovers in determining the extent of synaptic adjustment in mice.

### Regulating heterologous synapsis through attachment to the nuclear envelope?

We document dramatic differences between heterologous synapsis on the standalone and the fused X chromosomes. This holds true upon perturbation of DSBs and crossovers, and also in worms where those pathways are not perturbed (Fig 4). Structurally, the two karyotypes we examined differ by a single phosphodiester bond between the telomeres of chromosomes X and V. However, this change could have multiple mechanistic implications on synapsis.

One possible explanation for the difference is the ability of the fused chromosome to undergo synaptic adjustment in addition to fold-back synapsis. This scenario implies that synaptic adjustment in worms can fully align the pseudo-homologs, entailing axial compression or expansion by a of factor of two.

Alternatively, the difference in heterologous synapsis between the two karyotypes stems from increased propensity of the fused X chromosome to undergo fold-back synapsis. Since extension of synapsis is sequence-independent (MacQueen et al. 2005) and not rate-limiting for completion of synapsis (Rog and Dernburg 2015), it is likely that initiation of fold-back synapsis is impacted by fusion.

In worms, synapsis initiation between homologous chromosomes occurs at Pairing Centers (MacQueen et al. 2005; Rog and Dernburg 2015), and is regulated by local homology and by the attachment of chromosomes to force-generating machinery in the cytoplasm (Sato et al. 2009). Initiation of fold-back synapsis, as examined here, occurred in the absence of sequence homology or Pairing Center proteins. This unconventional initiation likely explains its delayed kinetics: it occurred much later in meiosis than synapsis initiation at homologous Pairing Centers (Fig S1 and MacQueen et al. 2002). The delayed kinetics likely reflects a higher energetic barrier that had to be overcome, and potentially a requirement for higher concentration of unassembled SC proteins (Harper et al. 2011).

However, the standalone X chromosomes we analyzed lacked not only the ability to assess local homology at the synapsis initiation site, but also attachment to force-generating machinery in the cytoplasm (Penkner et al. 2007; Sato et al. 2009). In addition to facilitating homology search (Wynne et al. 2012), dynein-driven chromosomal movements promote synapsis initiation (Sato et al. 2009). It is likely that the same effect could also promote initiation of fold-back synapsis when the folded-back chromosome is fused to an autosome. Pushing and pulling the chromosome may modify the axis—chemically or physically—which then primes it for association with the SC. A non-mutually exclusive hypothesis is that the proximity to a synapsed chromosome promotes fold-back synapsis, perhaps by increasing the local concentration of SC subunits.

### Outlook

Heterologous synapsis is a prevalent feature of the meiotic program (Zickler and Kleckner 1999), and may be as widespread evolutionarily as the SC itself. While heterologous synapsis has the potential to promote illegitimate recombination, an important future avenue of research will be to characterize various kinds of illegitimate crossovers (e.g., León-Ortiz et al. 2018), and the degree by which heterologous synapsis promotes them. In addition, recent work from our lab suggests another potential danger posed by heterologous synapsis: elevated meiotic sister-chromatid exchanges (Almanzar et al. 2021). Our finding of heterologous synapsis suppression by DSBs and crossovers opens the door to future studies of this potentially important pathway of preserving genome integrity during sexual reproduction.

## Methods

### Worm strains, growth conditions and quantification of brood sizes

Worms were maintained according to standard protocols (Brenner 1974). Auxin plates contained a final concentration of 1mM auxin (VWR Cat#AAA10556-36) prepared from a 500mM stock in ethanol (Zhang et al. 2015). For auxin depletion, only animals that grew on auxin from hatching were analyzed. For analysis of the pseudo-homologs, males lacking the fusion chromosome were mated to hermaphrodites homozygous for the *ypT27* fusion (both animals had identical genotypes otherwise). F1 cross-progeny hermaphrodites were analyzed. For brood counts, four age-matched L4s were picked onto individual NGM or auxin plates, and transferred onto new plates every 24 hours for three consecutive days. Adult progeny was counted on all plates and summed.

### CRISPR/Cas9 construction and injection

*aid∷cosa-1* repair template consisted of an *aid* degron in-frame with the open reading frame of *cosa-1* through a six-amino acid linker of glycines and serines, GSGSSG (Zhang et al. 2015). The PAM sequence CGG was mutated to CGC to prevent re-cutting. The template was ordered as two ultramers from IDT, with a 35 bases overlap between them (Paix et al. 2016). Guide RNA was also ordered from IDT (Fig S2). The resulting strain was verified through PCR and Sanger sequencing (see Reagent List for primer pairs). CRISPR/Cas9 injections were carried out on Zeiss Axio Vert.A1 using Eppendorf FemtoJet 4i and Eppendorf TransferMan 4r.

### Immunofluorescence staining

Worm gonad dissection and immunofluorescence staining were carried out following (Phillips et al. 2009) with a slightly modified version of egg buffer (25mM HEPES pH 7.3, 118mM NaCl, 48mM KCl, 2mM CaCl_2_, 2mM MgCl_2_; Zhang and Kuhn 2013). The primary antibodies used were: goat anti-SYP-1 (gift of Abby Dernburg; MacQueen et al. 2002), rabbit anti-SYP-5 (gift of Yumi Kim; Zhang et al. 2020), and guinea pig anti-HTP-3 (Hurlock et al. 2020). These were used at 1:400 dilution. Secondary antibodies include donkey anti-rabbit, donkey anti-guinea pig, and donkey anti-goat from Jackson ImmunoResearch, conjugated with either Alexa Fluor 647 or Alexa Fluor 488. These were used at 1:100 dilution.

### Imaging and image analysis

Confocal images were acquired through Zen Black 2.3 on a Zeiss LSM 880 equipped with an Airyscan and a Zeiss Plan-APOCHROME 63x/1.4 NA oil objective. Z-stacks were taken with 0.159μm intervals. Pixel size was 0.0353μm. Images were Airyscan-processed in Zen 2.3 Blue and then quantified through 3D reconstructions in Imaris 9.5.1 or 9.7.2 (Oxford Instruments), where they were resampled to 0.0353μm intervals along the z axis.

For DAPI body quantifications, adult hermaphrodites that were hatched and grown on auxin were dissected and stained with anti-HTP-3 antibodies and DAPI. DAPI bodies in diakinesis nuclei were counted in z-stack 3D reconstructions using Imaris. HTP-3 was used to help distinguish between bivalents and univalents.

For analysis of fused X chromosomes, age-matched *ypT27(+/−)* worms (24 or 48 hours post-L4) were grown on either NGM (for *dsb-2* and *spo-11(+)*) or auxin plates (for *spo-11(−)* and *cosa-1(kd)*) at 20◻ before dissection. After acquiring confocal z-stacks, maximum intensity projections were generated for each gonad. 2D analyses were carried out directly from the projections, scoring nuclei in pachytene. For 3D analyses, the projections were used to find the regions of interest (ROIs). ROI was defined as nuclei between the end of transition zone (based on DAPI staining; MacQueen et al. 2002; Colaiácovo et al. 2003; Harper et al. 2011) and the start of diplotene (first row of nuclei with discernable SC disassembly). Each nucleus within the ROI was marked. Then, in Imaris, 3D reconstructions were resampled along the z axis from 0.159μm to 0.0353μm to create a cubic voxel size, and if necessary, the channels were also aligned. Chromosomes lacking an SC (HTP-3-positive and SYP-1- or SYP-5-negative), were scored, and verified by tilting the nuclei. After scoring the nuclei, the ROI was divided into three bins of same size based on the number of rows. Fold-back quantification followed the same process, except lacking the *ypT27* fusion.

### Analysis of DSB perturbation in *dsb-2* animals

From previously published results in figure 1D by Rosu et al. 2013, we calculated the average number of crossovers in any given nucleus at 24 and 48 hours post-L4. The average number of DSBs was calculated from the number of crossovers using the equation in Yokoo et al. 2012.

### Calculation of R_S_ and R_F_

We calculated the rates of complete fold-back synapsis on the standalone X chromosomes (R_S_) and complete heterologous synapsis on the fused chromosomes (R_F_) by finding the best-fit to a binomial distribution. Binomial distributions for probabilities with 0.001 increments of rates were calculated for each condition, *i.e*. nuclei with 0, 1, or 2 asynapsed regions. R_S_ corresponds to the rate with the smallest sum of squares of difference to the observation. R_F_ was calculated by testing potential heterologous synapsis rates of the fused overhang that increased in increments of 0.001, with Rs for the other chromosome derived from the fold-back analysis above. Results were calculated against observed values, and R_F_ corresponds to the probability with the smallest sum of squares of differences. Averages of all gonads in each condition were used as the final R_S_ and R_F_. All calculations were conducted in Excel (Microsoft Corporation) using BINOM.DIST(number, 2, probability, false) function.

### Statistical Analysis

Raw data were compiled in Excel (Microsoft Corporation). All statistical analysis were performed in PAST 4 (Hammer et al. 2001), with the exception of goodness-of-fit chi-square test for binomial distribution, which was performed manually in Excel.

The statistical significance value α=0.05 was used for all analyses. In all figures, significant p values were designated as follow: (*) for 0.05≥p>0.005, (**) for 0.005≥p>0.0005, and (***) for p≤0.0005. Normality tests were performed across all datasets prior to further statistical analysis. Datasets where the majority of data was normally distributed were analyzed through Student’s t-test (equal variance) or Welch’s t-test (unequal variance); Mann-Whitney U test was used otherwise.

## Acknowledgement

We would like to thank Shawn Ahmed for worm strains, Yumi Kim and Abby Dernburg for antibodies, members of the Rog lab for discussions, Lexy Diezmann, Yuxuan Li, and Yifan Sun for statistical advice, Lisa Kursel and Yuval Mazor for critical reading of the manuscript and editorial suggestions, Jorgensen Lab for NGM and auxin plates, the Taft-Nicholson Center for a Summer Faculty residency, and The University of Utah Cell Imaging Core for access to Imaris software.

## Declarations

### Funding

Worm strains were provided by the CGC, which is funded by NIH Office of Research Infrastructure Programs (P40 OD010440). Work in the Rog lab is supported by R35GM128804 grant from NIGMS, and start-up funds from the University of Utah.

### Conflict of Interest

The authors declare no interest of conflict.

### Authors’ Contributions

HL carried out most experimental work. SGG performed the 2D analysis (Fig 3d) and made the initial observation regarding *spo-11(−)* animals. HL and OR analyzed the data and wrote the manuscript, with input from all authors.

---

**Supplementary Fig. 1.**
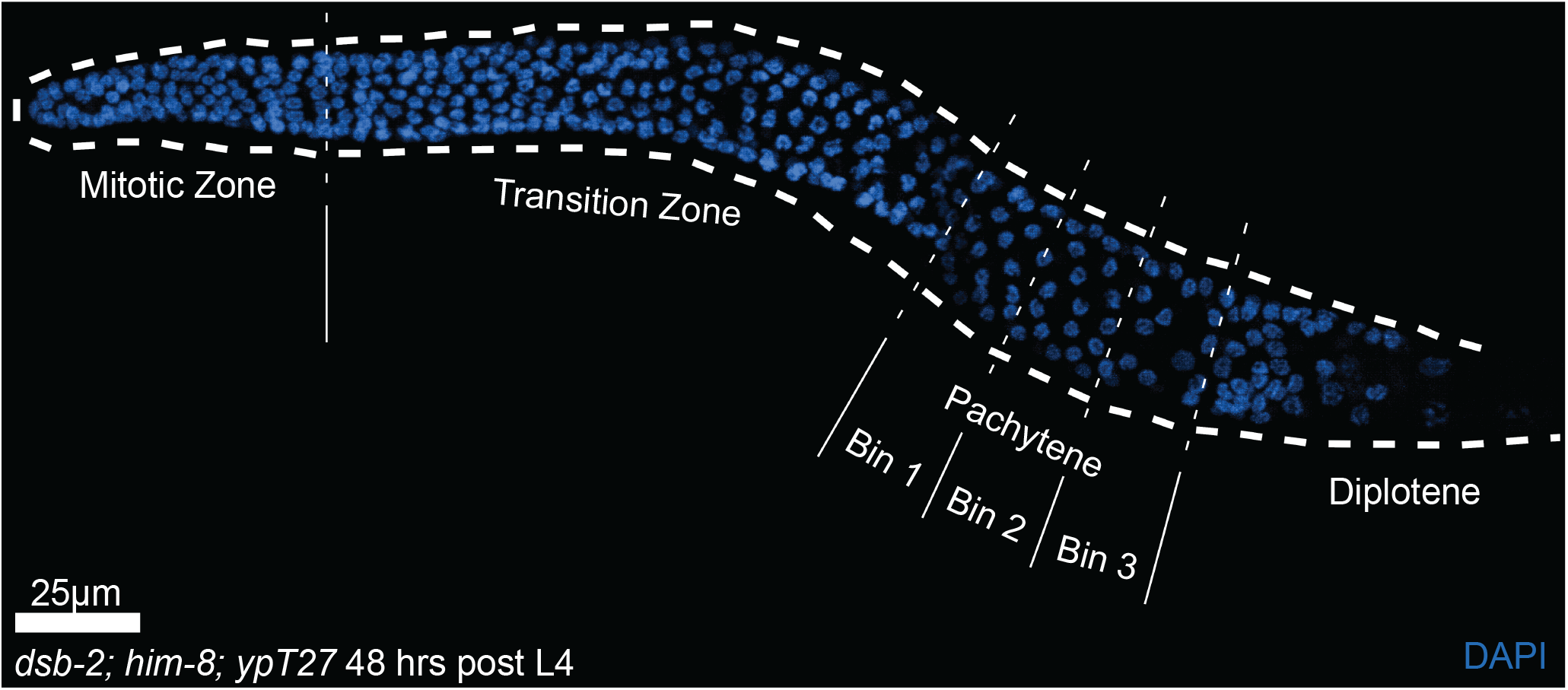
Experimental system to study heterologous synapsis in *C. elegans*. Overview of meiotic progression in the *C. elegans* gonad (*dsb-2; him-8*, 48 hours post-L4 gonad is shown). Dissected gonad is stained with DAPI to label DNA (blue). Meiosis progresses from left to right. The *him-8* mutation, present in all strains used in this study, extends the transition zone, which spans only ~10% of the gonad in *him-8(+)* animals (Harper *et al*. 2011). The pachytene region is divided into three bins of equal length in order to observe the temporal progression of heterologous synapsis.

**Supplementary Fig. 2.**
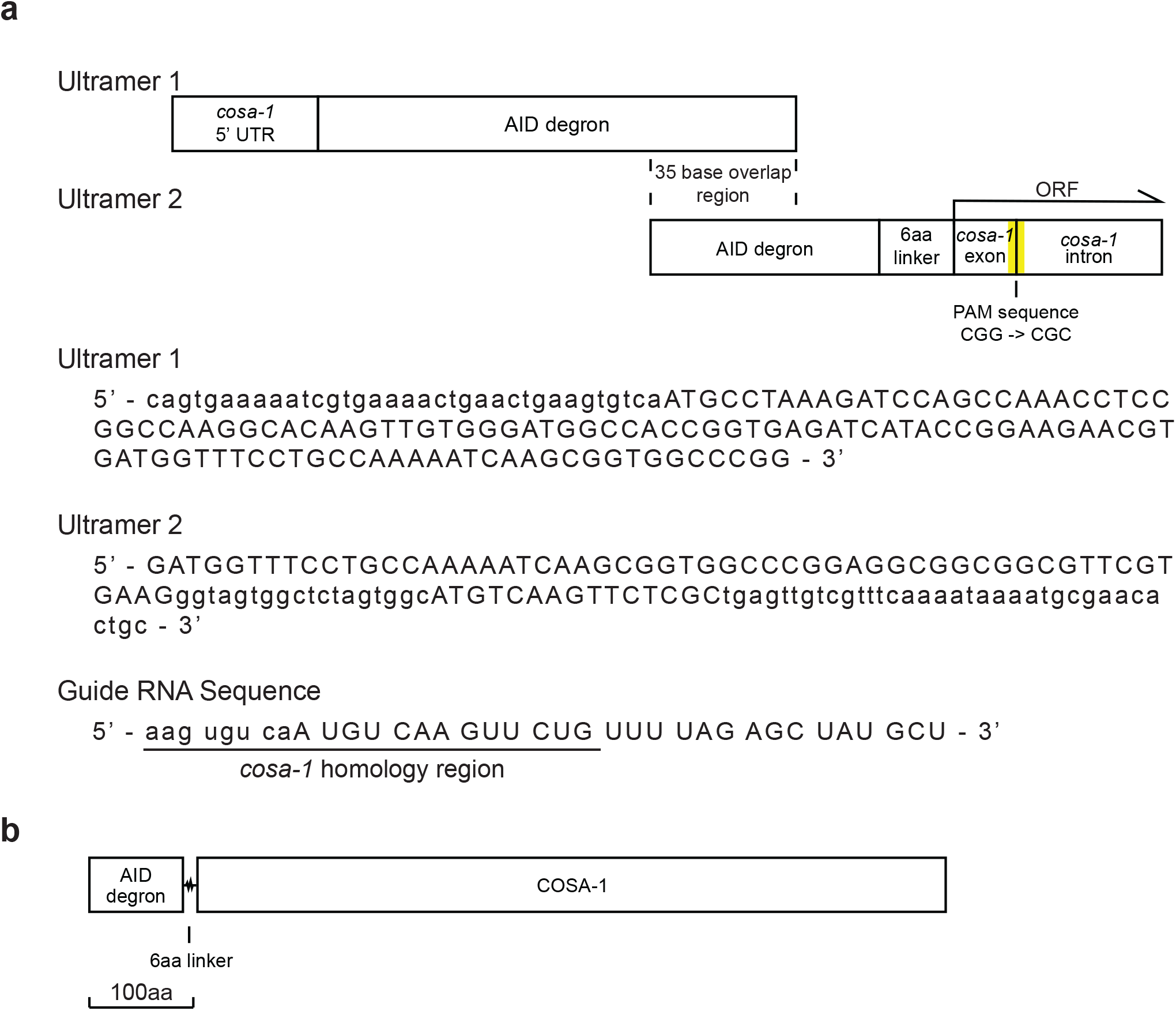
Construction of *aid∷COSA-1* using CRISPR/Cas9. A. Two single-stranded oligonucleotides serve as the template to create *aid∷COSA-1*. Top: Schematic of the ultramer templates. AID degron tag (Zhang *et al*. 2015) is inserted at the N-terminus of COSA-1. The two templates are 150 and 125 nucleotides long (ultramers, IDT), with a 35-nucleotide overlap. The template consists of homology to the *cosa-1* 5’ untranslated region, AID degron sequence, a six-amino acid linker of glycines and serines, and homology to the first exon and intron of *cosa-1*. The CGG PAM sequence (yellow), at the end of the first *cosa-1* exon and the beginning of the first *cosa-1* intron, is mutated to CGC to avoid re-cutting. Middle: the ultramer sequences. Bottom: the guide RNA sequence. Exon sequences are shown in uppercase. B. Schematic of the resulting AID∷COSA-1 protein, drawn to scale.

**Supplementary Fig. 3.**
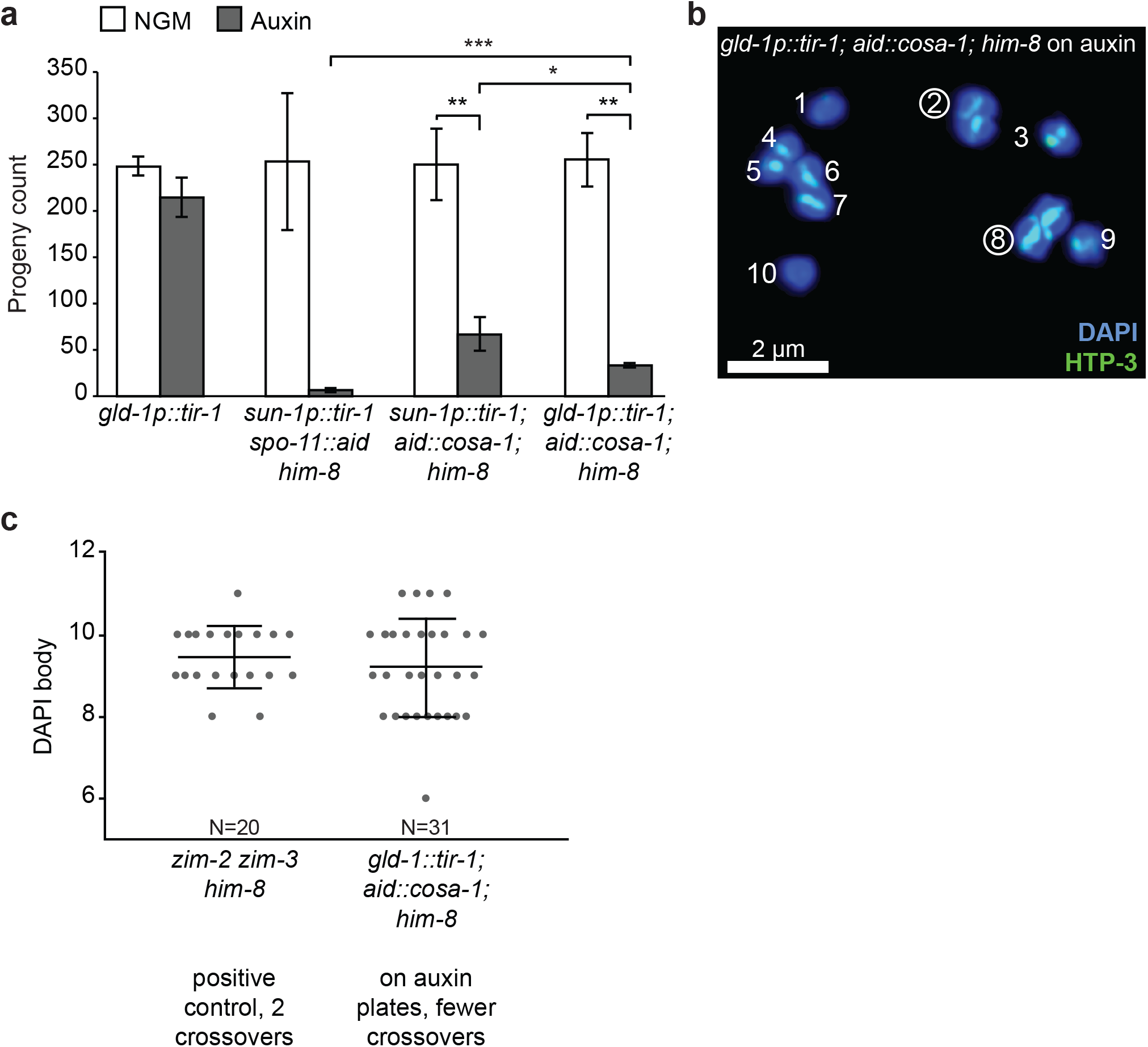
Optimization and characterization of AID∷COSA-1. A. *gld-1* promoter is more robust than *sun-1* promoter in driving AID∷COSA-1 degradation. Average self-progeny is shown in the presence and absence of auxin. *gld-1p∷tir-1* is used as the negative control while *sun-1p∷tir-1 spo-11∷aid him-8* (i.e. *spo-11(−)*) is used as a positive control. TIR-1 driven by either the *sun-1* or *gld-1* promoters degrades AID∷COSA-1 when on auxin, yielding a lower brood size (p<0.005 compared to on NGM plates, Welch’s t-test). However, TIR-1 driven by *gld-1* promotor is more robust (p=0.0475 compared with *sun-1* promotor, Welch’s t-test). N represents the number of worms assessed. Genotypes are indicated at the bottom. B. Degradation of AID∷COSA-1 on auxin by *gld-1* promoter-driven TIR-1 yields more than six DAPI bodies in diakinesis. Numbers indicate counted DAPI bodies. Circled numbers indicate linked homologs (bivalents). Blue, DAPI bodies. Green, HTP-3 (axis). Scale bar=2 μm. C. *gld-1p∷tir-1; aid∷COSA-1; him-8* yields similar number of DAPI bodies as *zim-2 zim-3 him-8* (p=0.3976, Student’s t-test), indicating the presence of about two crossovers per meiosis (Phillips and Dernburg 2006). N represents the number of diakinesis nuclei assessed, each represented by a dot. Bars show mean ± SD. Genotypes and their effects are indicated at the bottom.

## Notes

### Competing Interest Statement

The authors have declared no competing interest.

